# Vps60 initiates formation of alternative membrane-bound ESCRT-III filaments

**DOI:** 10.1101/2022.02.16.480668

**Authors:** Anna-Katharina Pfitzner, Henry Zivkovic, Frédéric Humbert, Aurélien Roux

## Abstract

Endosomal sorting complex required for transport-III (ESCRT-III)-driven membrane remodeling participates in many crucial cellular functions, from cell division to endosome maturation, and occurs on essentially all cellular organelles. In eukaryotes, ESCRT-III displays a remarkable molecular diversity in its subunits which may have been acquired through evolution to perform novel cellular functions. Here, we describe and characterize a novel ESCRT-III polymer initiated by the subunit Vps60. Membrane-bound Vps60 polymers recruit ESCRT-III subunits Vps2, Vps24, Did2 and Ist1, and undergo polymer turnover powered by the ATPase Vps4. Snf7- and Vps60 filaments can coexist on membranes without interacting. Their nucleation, polymerization and recruitment of downstream subunits remains unaffected by the presence of the respective other polymer. Taken together, our results suggest Vps60 and Snf7 form distinct ESCRT-III polymers, which overall, supports the notion of evolutionary diversification of ESCRT-III assemblies to perform specific cellular functions.

## Introduction

Lipid membranes are a hallmark of living cells. To maintain functionality, they require constant remodeling by dedicated machineries like the Endosomal Sorting Complex Required for Transport-III (ESCRT-III). Presumed to be the first membrane remodeling machinery to have evolved^1,2^, ESCRT-III acts on virtually all cellular membranes to promote membrane fission from within membrane necks, a process that is essential for many cellular functions such as formation of intralumenal vesicles (ILVs) from endosomal membranes, cytokinetic abscission of the plasma membrane, reformation of the nuclear envelope, and closure of autophagosomes^3–5^. Moreover, ESCRT-III catalyzes budding of various virions in eukaryotes and archaea^6–11^ and functions in repairing lipid membranes, as shown for the eukaryotic and bacterial plasma, lysosomal, nuclear and plastid membranes^2,12–18^, a function that is essential to sustain vacuolar confinement of pathogens during certain infections^19,20^. Unlike other membrane remodeling machineries, ESCRT-III can also function with a reverse orientation promoting membrane fission from the outside of membrane necks during release of peroxisomes, recycling of endosomes and lipid droplet formation^21–23^. Despite its ubiquitous role in vital functions, the mechanism by which ESCRT-III performs membrane remodeling and the machinery’s adaptation to its various cellular functions is not fully understood.

Canonically recruited to endosomal membranes by ESCRT-II, ESCRT-III assembly starts with Vps20 (CHMP6) followed by subunits Snf7 (CHMP4B), Vps2 (CHMP2A) and Vps24 (CHMP3) before being likely completed by subunits Did2 (CHMP1B) and Ist1 (IST1). Besides the canonical pathway, other nucleators such as Bro1 (ALIX) and Chm7 (CHMP7) can recruit ESCRT-III to diverse cellular membranes^24–29^. Vps2 and Vps24 as well as Vps2 and Did2 bind Snf7 synergistically and then recruit the AAA-ATPase Vps4^30–36^, which induces subunit turnover within ESCRT-III polymers promoting either disassembly^37–39^, growth^40^ or sequential subunit polymerization^41^. In cells, Vps4-dependent polymer remodeling is indispensable for ESCRT-III function^40,42,43^. Upon recruitment, ESCRT-III subunits assemble into filaments with diverse stoichiometries and shapes ranging from spirals^44–47^ to tubular helices^37,48–51^ and spiraling membrane tubes^52,53^. Sequential succession of these various ESCRT-III filaments has recently been suggested to promote ESCRT-III mediated membrane remodeling^54,55^.

Aside from the well characterized core subunits Snf7, Vps2 and Vps24, several accessory ESCRT-III subunits have been identified based on a deletion phenotype indicative of disturbed ILVs formation^56^ and their secondary structure organization^57^, which is highly conserved among ESCRT-III proteins even across species. As one of those accessory subunits, the function of Vps60/Mos10 (CHMP5), though briefly associated with ESCRT-III disassembly^58,59^, remains poorly understood to this day. A recent analysis of genetic interactions between ESCRT-III subunits, however, places Vps60 more central in an interaction network^36^ implying potentially a more important function for Vps60 than previously recognized. We thus decided to perform a functional characterization of Vps60 and its interactions with other ESCRT-III subunits as well as the ATPase Vps4.

## Results

### Vps60 behaves like an early ESCRT-III protein

We here set out to characterize the function of ESCRT-III accessory subunit Vps60. In general, most ESCRT-III subunits or submodules, though to varying degrees, can polymerize into membrane-bound filaments which often depict preferential binding to a specific membrane curvature range^53,60,61^. Snf7, the initial ESCRT-III subunit, polymerized spontaneously (Fig. 1A) on giant unilamellar vesicles (GUV), whereas downstream submodules Vps2-Vps24 (Fig. 1C)^61,62^ and Vps2-Did2-Ist1 (Fig. 1D)^41^ required activation, here provided by an acidic buffer (Fig. 1A), to polymerize. Interestingly, Atto565-Vps60 bound spontaneously to GUVs (Fig. 1B) and was, similar to Snf7, recruited efficiently to flat non-deformable supported lipid bilayers (SLBs) (Fig. S1A-B).

**Figure 1.**
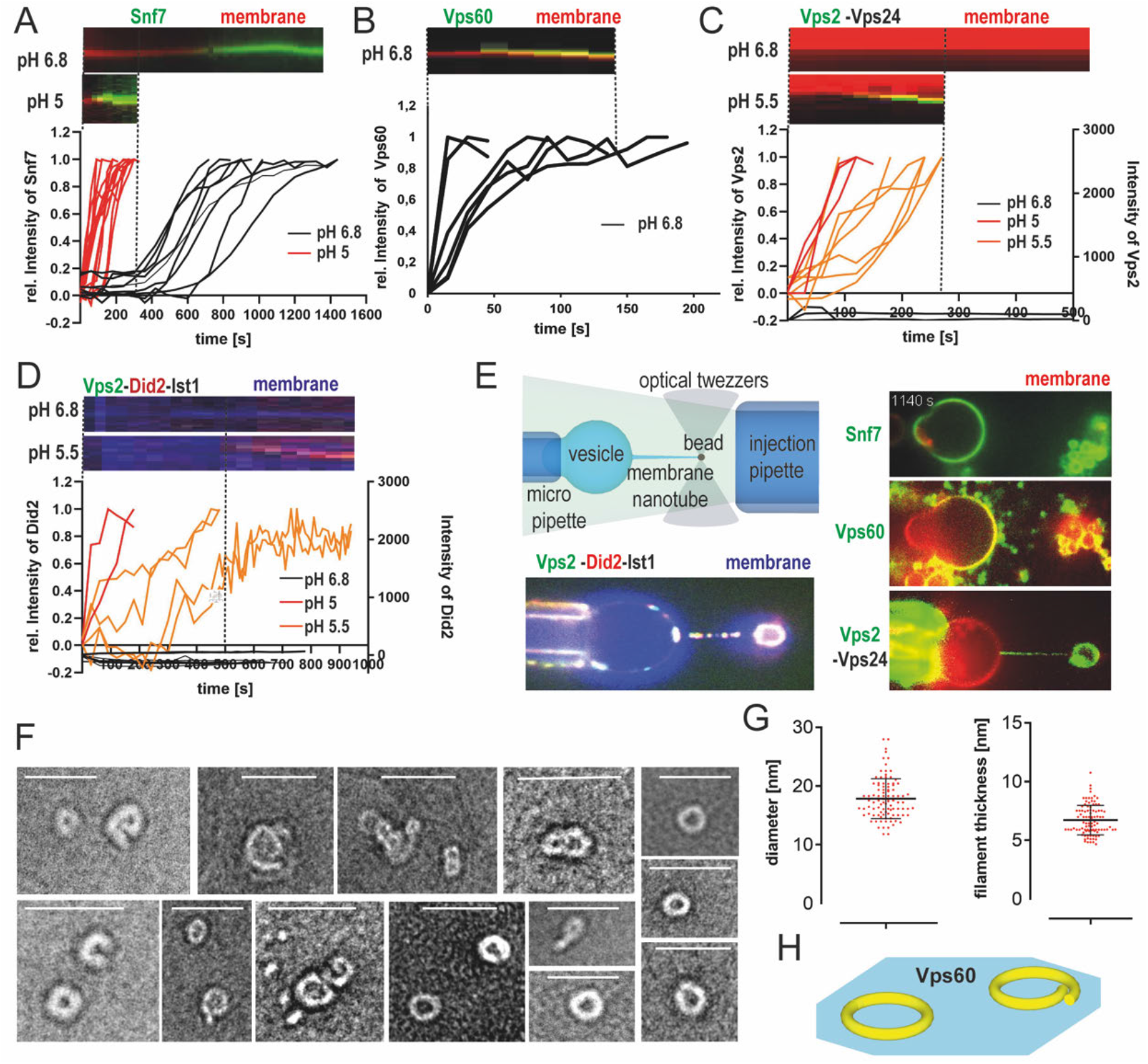
Comparison of biochemical properties of ESCRT-III proteins. A-D. Quantification and kymographs of dynamics of Snf7 (A), Vps60 (B), Vps2-Vps24 (C) or Vps2-Did2-Ist1 (D) binding to membrane at indicated pH. E. Schematic representation of membrane nanotube pulling and confocal microscopy images of Snf7-Alex488 (green), Vps60 (green), Vps2-Alex488 (green) and Vps24 or Vps2-Alex488 (green) and Did2-Atto565 (red) and Ist1 binding to membrane nanotubes (red, blue). F. Negative stain electron micrographs of Vps60 filaments polymerized on LUVs (scale bar: 100nm). G. quantification of experiment described in F. H. Schematic representation of Vps60 filaments.

After establishing Vps60’s affinity for membrane, we next asked whether the protein depicts a membrane curvature-preference. To this end, we injected labeled subunits in the vicinity of membrane nanotubes, the latter made by pulling beads adhered to GUVs with optical tweezers (see Methods and Fig. 1E). The described set-up produces highly curved and flat membranes close to each other, allowing us to evaluate curvature-dependent binding in a wide range. As previously reported, Snf7 bound exclusively flat membrane (Fig. 1E, S1C)^61^. Likewise, Vps60 is strongly recruited along the GUV’s flat membrane, whereas only minor binding is observed along the highly curved nanotube (Fig. 1E, S1E). In contrast, both downstream ESCRT-III submodules, Vps2-Vps24 and Vps2-Did2-Ist1, bind predominantly to highly curved nanotubes (Fig. 1E, S1D, F).

Upon membrane binding, all characterized ESCRT-III subunits polymerize into filaments^44–47,49,52,63,64^. To test if Vps60 behaves likewise, we performed negative stain electron microscopy of large unilamellar vesicles (LUVs) incubated with Vps60. Indeed, Vps60 formed ring-shaped filaments with an average diameter of 18.9±3.4 nm (Fig. 1F-H) similar to ring filaments described for Snf7^60,65^. In disagreement with our findings, a recent study reported Vps60 to form wide ranging spirals reminiscent to the Snf7 ones^66^. This discrepancy might arise from different experimental conditions as Banjade and colleagues used higher protein concentration which might help propagating spiral growth. Besides rings, we observed polymers with one inward-curled tip as it would be expected for a spiral initiator (Fig. 1F, H). Snf7 spiral polymers were previously suggested to grow out of ring-shaped filaments upon their spontaneous breakage^60,67^. In analogy, curled Vps60 polymers might arise from rupture of ring filaments. The lack of large spirals may potentially be due to filament-specific properties that control its polymerization rate such as filament thickness which is higher for Vps60-(6.7 ±1.3 nm, Fig. 1G) than for Snf7-filaments (5 nm) ^65,68^. Alternatively, curled filaments might result from ring breakage during sample preparation. We occasionally observed filaments which resemble stacks of rings (Fig. 1F), which might arise from buckling of curled filaments similar to buckling of other ESCRT-III polymers^52,65^.

In summary, Vps60 depicts characteristics similar to the early ESCRT-III protein Snf7 and clearly distinct from the properties of downstream modules Vps2-Vps24 and Vps2-Did2-Ist1. Curvature preference and filament shape suggest that Vps60 polymers potentially form a membrane binding interface perpendicular to the helical axis like Snf7-Vps2-Vps24 and, presumably, Snf7 polymers^52,69^ (Fig. 1 H). The higher curvature binding preference of later subunits (Vps2-Vps24, Did2-Ist1) cohere with their binding interface parallel to the helical axis^49,50,63,64^ (Fig. 1E).

### Spontaneous nucleation of Vps60 on membrane is highly efficient

As both, Vps60 and Snf7, polymerize spontaneously on membrane, we next set out to compare their nucleation capacities. We therefore analyzed the nucleation rate of Atto565-Vps60 on SLBs (Fig. 2A-B). Below 50 nM, Atto565-Vps60 nucleation events increased linearly with respect to protein concentration, whereas above 50 nM, their number seemingly increased exponentially. In contrast to the estimated nucleation rate, which was likely underestimated due to overlapping Vps60 puncta, images at concentrations between 150 nM and 250 nM do not suggest a saturation of Vps60 binding. These results point towards Vps60 displaying a higher intrinsic nucleation rate on membrane than Snf7, which, does not spontaneously nucleate below a concentration of 300 nM. Vps60 however does not form growing patches like Snf7, but instead binding manifests itself in accumulation of puncta (Fig. 2C) as well as an overall increase of intensity on the membrane. Growth of Snf7 patches was previously explained by breaking of preexisting spirals into multiple smaller spirals from which protein polymerization could continue^60^. Filament breaking thus fuels a chain reaction from a single nucleation event, leading to expanding protein patches formed of hundreds of growing spirals. Observed filament structures of Vps60 (Fig. 1C) suggested that filament breaking might occur less frequently and that spiral growth was partially or completely inhibited, explaining why no growing Vps60-patches were observed (Fig. 2C).

**Figure 2.**
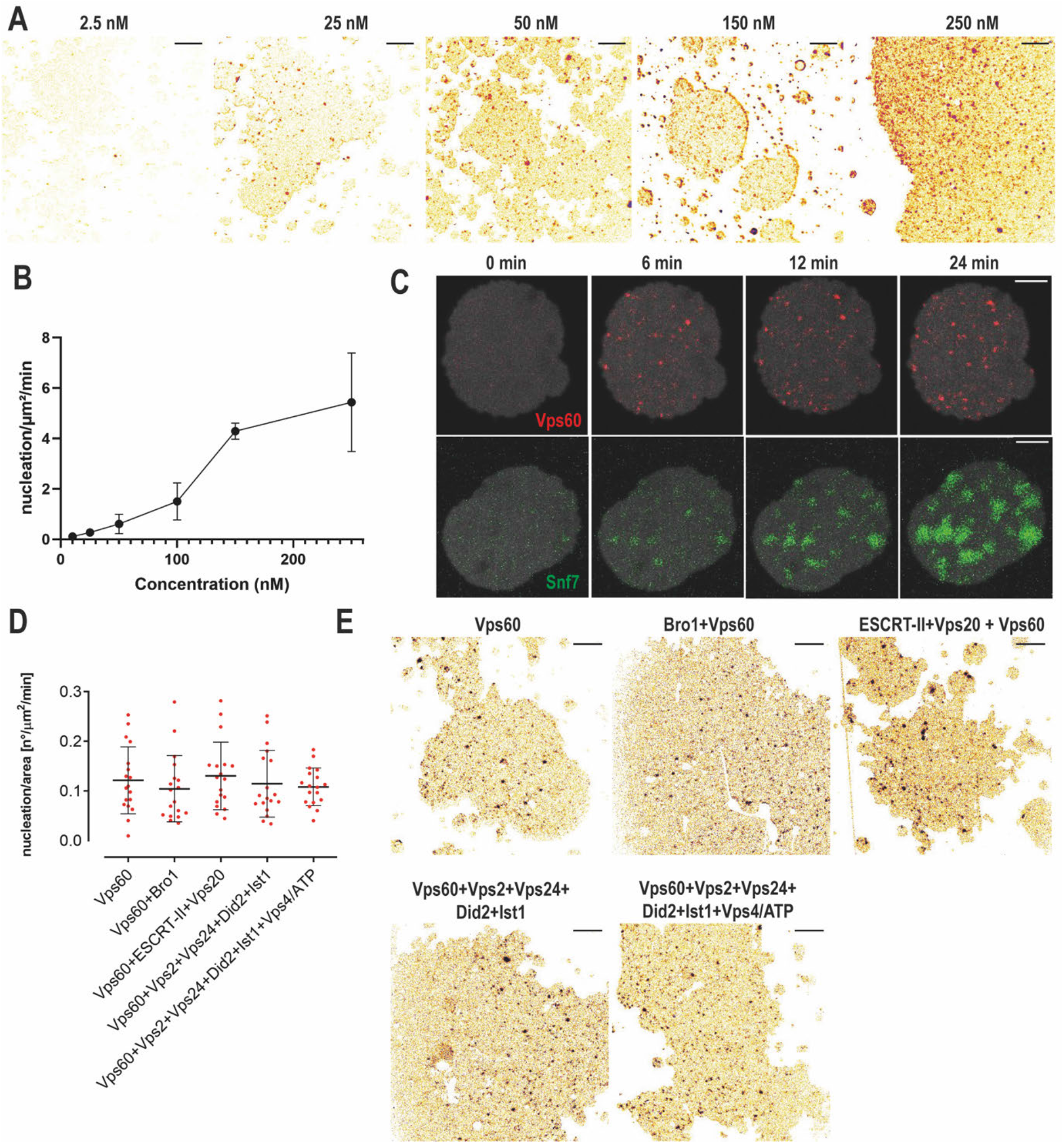
Nucleation properties of Vps60 polymerization on membrane. A. Confocal images of SLBs incubated with indicated concentration of Vps60 (scale bar 10 μm). B. Quantification of experiment described in A. C. Timelapse experiments of SLBs (gray) incubated with Vps60 (red, upper panel) or Snf7 (green, lower panel). D, E. Quantification (D) and confocal images (E) of SLBs incubated with Vps60 and the indicated proteins

Spontaneous nucleation rates for Snf7 (Fig. S2C-D)^60,70,71^ can be increased by dedicated ESCRT-III nucleators, like Bro1 and the ESCRT-II-Vps20 complex^26,65,72^. As Vps60 encompasses a Bro1-interaction domain and as its human homologue CHMP5 binds to the Bro1-domain containing protein Brox^73^, we wondered if Vps60 is targeted by these nucleators. Addition of Bro1 or ESCRT-II-Vps20 did not increase the Vps60 nucleation rate (Fig. 2D-E) nor the overall amount of protein recruited to membrane (Fig. S2A), indicating that Vps60 is not nucleated by these proteins. Vps60 could, however, interact with other ESCRT-III nucleators like Chm7^36^ or have other, Vps60-specific nucleators. As Snf7 membrane recruitment is strongly inhibited by downstream ESCRT-III proteins ^68^, we decided to also monitor Vps60 nucleation rates in presence of Vps2, Vps24, Did2, Ist1 and the ATPase Vps4. However, we did not observe any effect on Vps60’s membrane binding regarding nor its nucleation rate (Fig. 2D-E) or its kinetics (Fig. S2B).

### Vps60 and Snf7 display mutually exclusive membrane binding patterns

To next address if Vps60 and Snf7 interact upon each other’s polymerization on membrane, we analyzed Vps60’s nucleation on SLBs preincubated with Snf7, both simultaneously polymerizing and in absence of Snf7. This revealed Vps60 binding is unaffected by the presence of Snf7 (Fig. 3A-B, S3A-B). Similarly, Snf7 patches grew normally on SLBs preincubated with Vps60 (Fig. S3C). In fact, no colocalization of Snf7 and Vps60 was observed, even when the whole membrane surface was covered, indicating that Vps60- and Snf7 membrane binding are mutually exclusive (Fig. 3C-D). Moreover, incubation of Snf7-patches with high concentrations of Vps60 resulted in a decrease in Snf7 intensity on the membrane and vice versa (Fig. S3D-G). This suggestsVps60 and Snf7 compete to bind on available membrane surface.

**Figure 3.**
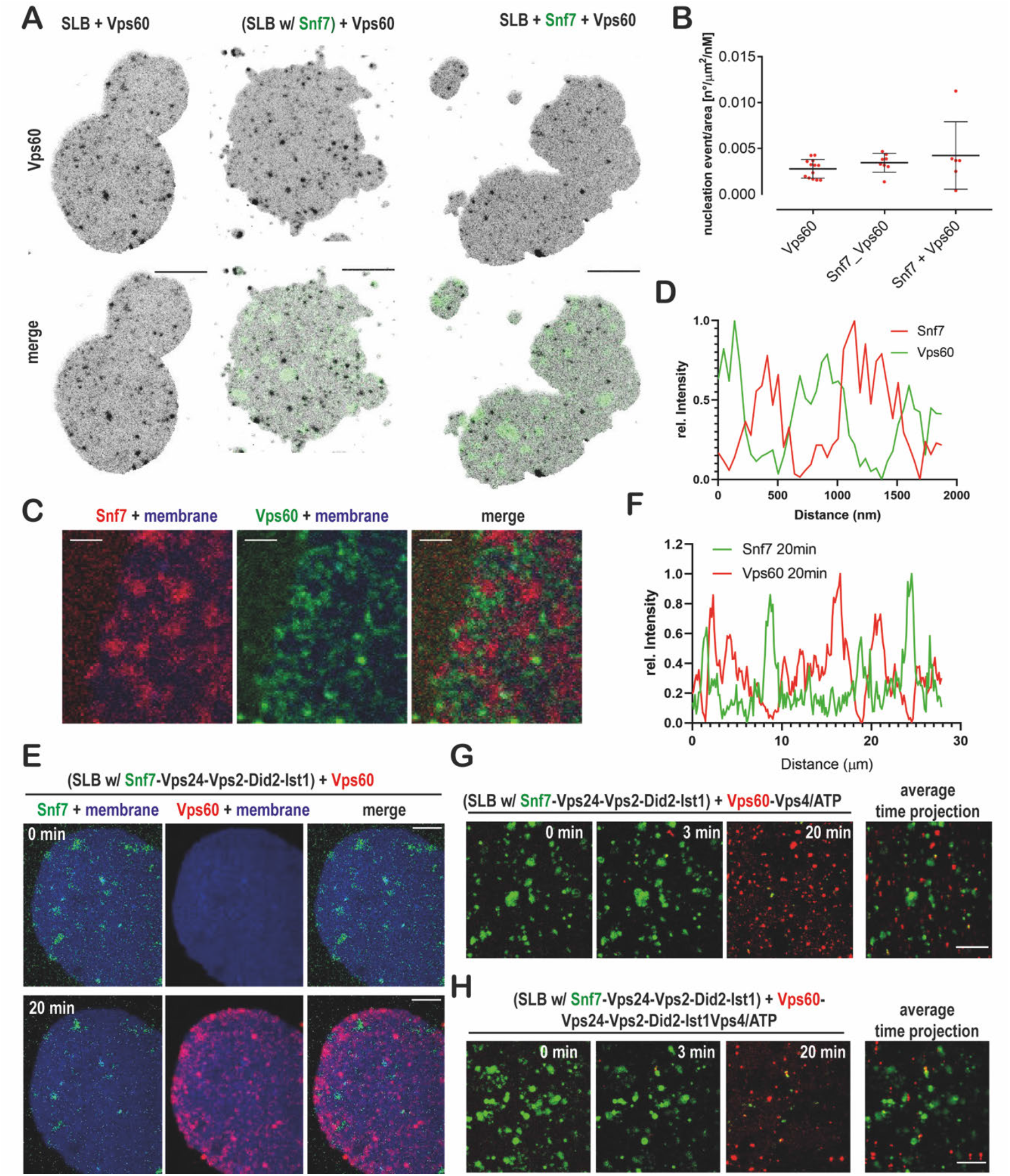
Vps60 and Snf7 bind membrane mutually exclusively. A. Confocal images of SLBs incubated with Vps60 (black) and Snf7 (green) where indicated (scale bar 10 μm). B. Quantification of experiments described in A. C. Confocal images of SLBs (blue) incubated with Snf7 (red) and Vps60 (green) (scale bar 2 μm). D. Plot of fluorescence profile of an exemplary membrane section from experiments described in C. D. Confocal images of timelapse experiment of addition of Vps60 (red) to SLBs (blue) pre-incubated with Snf7 (green), Vps2, Vps24, Did2 and Ist1. F. Plot of fluorescence profile of an exemplary membrane section (t =20 min) from experiments described in F.

Following polymerization, Snf7 filaments recruit downstream subunits starting with Vps2-Vps24^68^, followed by Vps2-Did2 and finally Ist1^41^. As Vps60 and Snf7 polymers co-exist on membrane without observable interaction, we asked if Vp60 is recruited into Snf7-based polymers by downstream subunits. We saw no recruitment of Vps60 to Snf7 polymers nor in the presence of Vps2, Vps24, Did2 and Ist1 or when Snf7-patches were pre-incubated with all the downstream ESCRT-III subunits (Fig. 3E-F). Instead, Vps60 bound membrane identically in the absence or presence of any downstream subunits. Likewise, no integration of Vps60 into Snf7-polymers was seen upon Vps4-induced filament turn-over, even in the presence of downstream subunits (Fig. 3G-H, S3FG).

In summary, we find Vps60- and Snf7 polymers to co-exist independently on membranes, and no recruitment of Vps60 to Snf7-based polymers. Altogether, with its characteristics similar to Snf7, we wondered if Vps60 might function parallelly to Snf7 as an alternative initiating subunit for a multi-subunit ESCRT-III filament.

### Vps60 polymers recruit downstream ESCRT-III subunits

To test our hypothesis, we studied the ESCRT-III subunit binding to SLBs pre-incubated with Vps60. Indeed, Alexa488-Vps2 was recruited strongly to Atto565-Vps60-covered SLBs in presence of Vps24 and Did2 (Fig. 4A-B). Similarly, we observe Vps60-mediated membrane binding of Alexa488-Vps24 in presence of Did2 and Vps2 (Fig. 4. C-D) and recruitment of Alexa488-Did2 when supplemented with Vps2 and Vps24 (Fig. 4 E, S4A). While Vps2-Vps24 are recruited to Snf7 polymers, Vps2-Vps24 was not sufficient for binding to Vps60-covered SLBs. Interestingly, we observed mild recruitment of Vps2-Did2 (Fig. 4A, E), indicating that Vps2-Did2 might act as major a recruitment complex in Vps60-based polymers, taking over the role of Vps2-Vps24 in Snf7-based polymers^62,68^. Vps24, however greatly increased Vps2-Did2 binding efficiency to Vps60-covered membranes, indicating that Vps24 might promote Vps2-Did2 heterofilament formation. As Vps60 and Did2 are reported to interact with Vps4-cofactor Vta1 ^58,59,74–77^, we tested if Vta1 affects the recruitment of Vps2-Did2-Vps24 to Vps60, which, however was not the case (Fig. S4C-D). Overall, we do not observe strong colocalization of Vps2, Vps24 and Did2 with Vps60 puncta, but find that their binding is specifically enhanced in the presence of Vps60 puncta. As Vps60-filaments do not form patches, it may be that Vps60 serves as a nucleation template for ESCRT-III polymers which can then diffuse along membranes.

**Figure 4.**
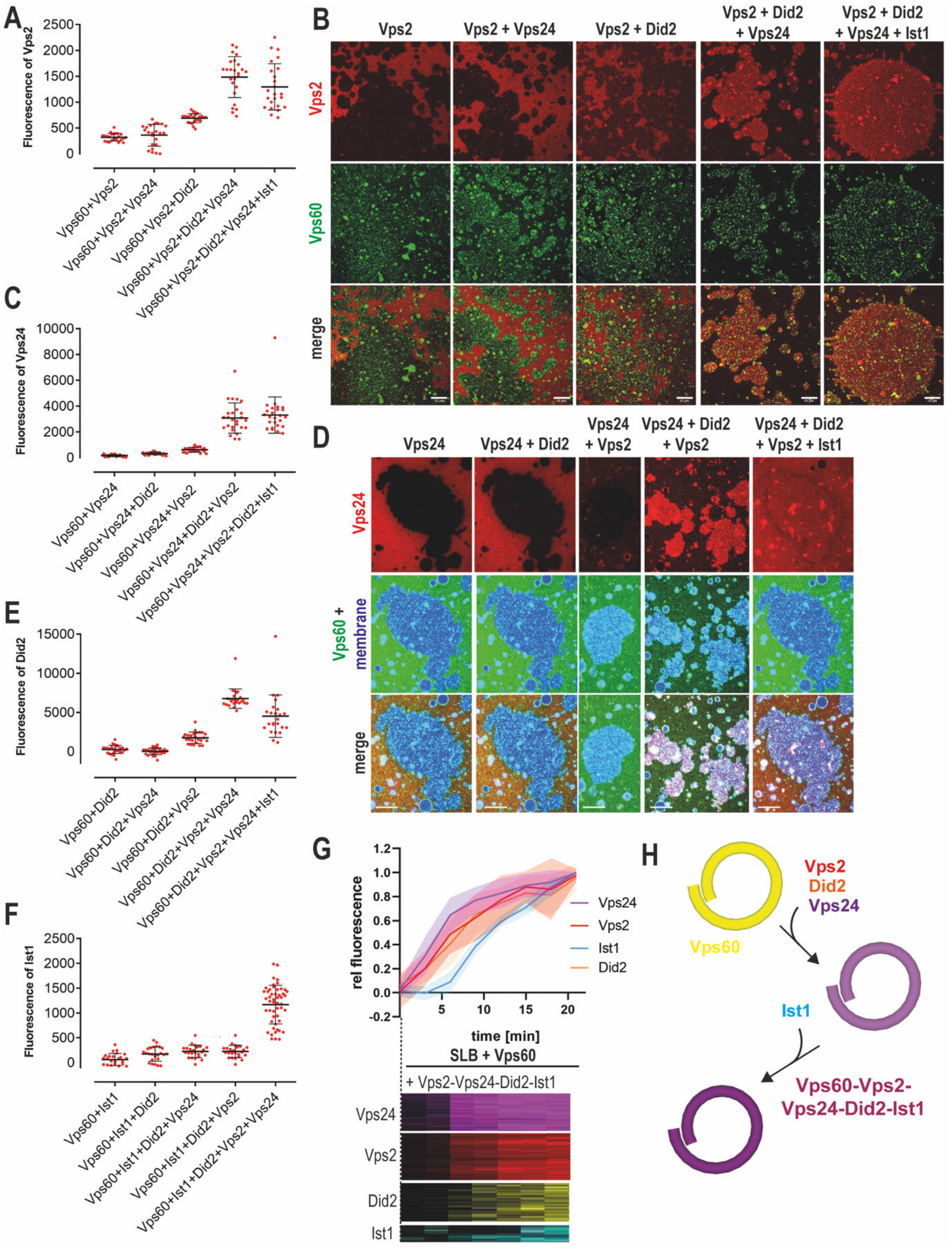
Vps60 recruit ESCRT-III subunits to membrane. A-B. Quantification of Vps2 fluorescence (A) and confocal images (B) of Vps60-covered SLBs incubated with Alexa488-Vps2 and the indicated protein mixture. C-D. Quantification of Vps24 fluorescence (C) and confocal images (D) of Vps60-covered SLBs incubated with Alexa488-Vps24 and the indicated protein mixture. E. Quantification of Did2 fluorescence of Vps60-covered SLBs incubated with Alexa488-Vps24 and the indicated protein mixture. F. Quantification of Ist1 fluorescence of Vps60-covered SLBs incubated with Alexa488-Vps24 and the indicated protein mixture. G. Kymographs and quantification of timelapse-experiments of Vps60-covered SLBs incubated with Alexa488-Vps24 (purple), Alexa488-Vps2 (red), Alexa488-Did2 (yellow) and Alexa488-Ist1 (cyan). H. Schematic representation of model for sequential binding of ESCRT-III subunits to Vps60-polymers.

Alexa488-Ist1, analogous to the other ESCRT-III subunits, was specifically recruited to Vps60-covered vesicles in presence of Vps2, Vps24 and Did2 (Fig. 4E, S4B). For this experiment, we used GUVs instead of SLBs as Ist1 formed aggregates in solution which sedimented on the SLBs precluding the monitoring of Vps60-induced binding. Ist1 binding pattern mirrored Did2 recruitment, which is consistent with the previously reported Did2-Ist1 heterodimer formation^78^ and suggests Ist1 incorporation into ESCRT-III polymers to only rely on this interaction with Did2.

During Snf7-mediated recruitment of downstream subunits, distinct recruitment kinetics can be observed, we thus asked if a similar temporal organization can be seen for Vps60-induced membrane binding. Timelapse imaging of Vps60-covered SLBs incubated with Vps2, Vps24, Did2 and Ist1 revealed a synchronic increase of Vps2-, Vps24- and Did2 intensity and a slightly delayed increase in Ist1 intensity (Fig. 4G). Overall, this result supports the notion of Vps2, Did2 and Vps24 together forming the initial recruitment complex (Fig. 4H). Subsequently, membrane-bound Did2 then triggers binding of Ist1 to ESCRT-III polymers. Compared to Snf7-mediated recruitment of Vps2-Vps24, binding of downstream subunits is slower when initiated by Vps60. Indeed, recruitment kinetics of Vps2-Vps24-Did2 to Vps60-filaments appears equivalent to Vps2-Did2 binding to Snf7-polymers^41^ supporting the notion that in Vps60-mediated assemblies, the two initial waves of subunits (Vps2-Vps24, and then Vps2-Did2) are condensed into a single Vps2-Vps24-Did2 wave.

### Vps60-nucleated ESCRT-III polymers undergo Vps4-mediated turnover

Previous studies demonstrated that ESCRT-III function in cells crucially depends on the ATPase activity of Vps4, which triggers filament turnover or remodeling^68,79,80^. To study if the Vps60-based polymers are likewise remodeled by Vps4, we performed timelapse imaging of SLBs pre-incubated with Vps60, Vps2, Vp24, Did2. Upon addition of Ist1, Vps4 and ATP, Vps2 and Vps24 intensities decreased rapidly, indicating subunit disassembly (Fig. 5A). In contrast, Did2 and Ist1 remained stably bound to membranes when Ist1 was in excess, whereas Vps4-triggered disassembly occurs in absence of Ist1 (Fig. 5A, S5A-B). These results suggest that Did2 is protected from disassembly by competitive binding of Ist1, and Vps4 to Did2 (Fig. S5E), a mechanism that we previously proposed for Snf7-based filaments^41^. As bound Ist1 itself undergoes continuous Vps4-triggred turnover, equilibrium between free Ist1 and Vps4 probably determines if Did2 is disassembled or stabilized.

**Figure 5.**
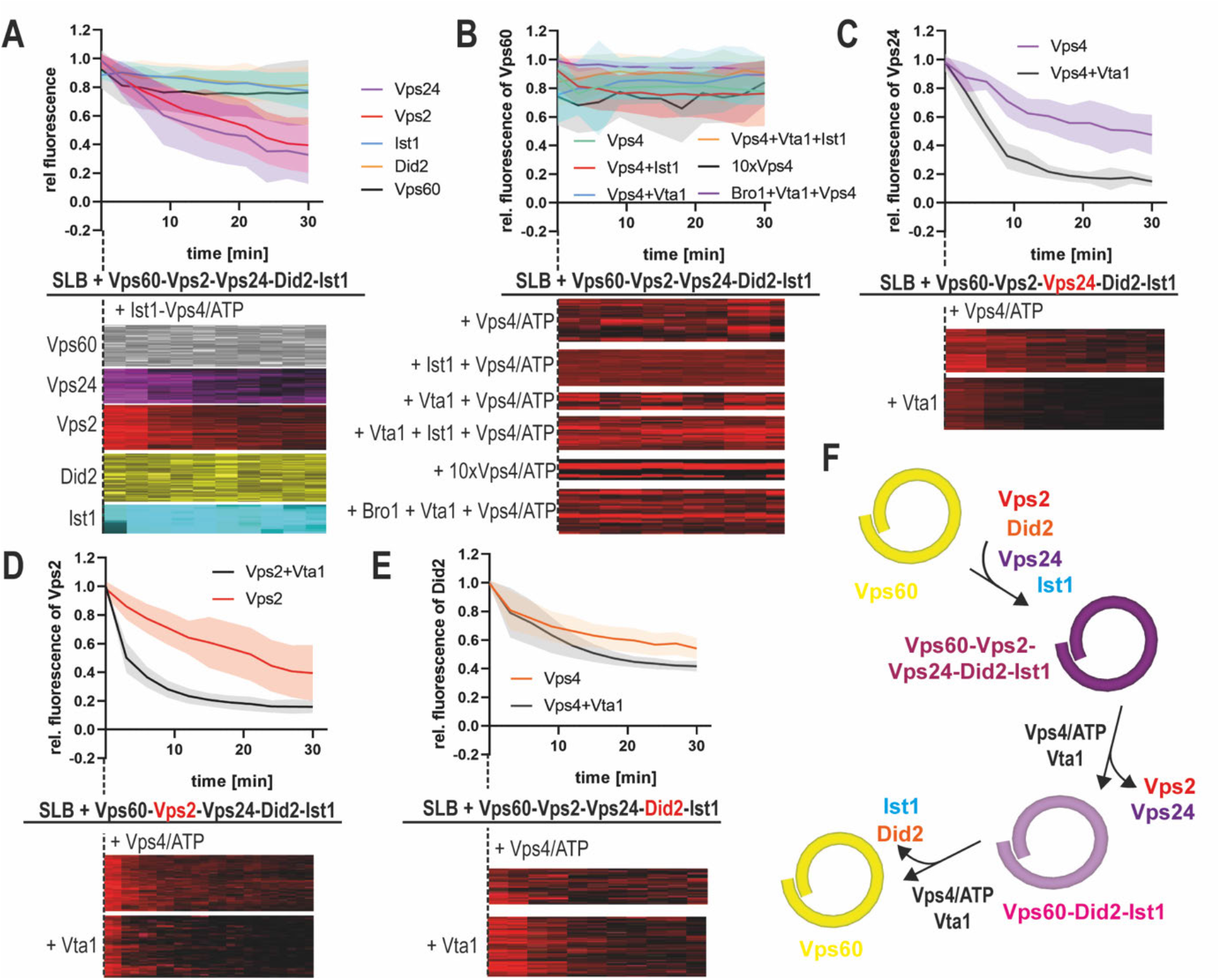
Vps4 triggers turnover of Vps60-based ESCRT-III polymers. A. Quantification of fluorescence intensities of indicated subunit and kymographs of timelapse experiments of addition of Ist1 and Vps4/ATP to SLBs pre-incubated with Vps60, Vps2, Vps24, Did2 and Ist1. B. Quantification of fluorescence intensities of Vps60 and kymographs of timelapse experiments of addition of Vps4/ATP and the indicated proteins to SLBs pre-incubated with Vps60, Vps2, Vps24, Did2 and Ist1. C. Quantification of Vps24 fluorescence intensities and kymographs of timelapse experiments of addition of Vps4/ATP and Vta1 where indicated to SLBs pre-incubated with Vps60, Vps2, Alexa488-Vps24, Did2 and Ist1. D. Quantification of Vps2 fluorescence intensities and kymographs of timelapse experiments of addition of Vps4/ATP and Vta1 where indicated to SLBs pre-incubated with Vps60, Vps24, Alexa488-Vps2, Did2 and Ist1. E. Quantification of Did2 fluorescence intensities and kymographs of timelapse experiments of addition of Vps4/ATP and Vta1 where indicated to SLBs pre-incubated with Vps60, Vps2, Vps24, Alexa488-Did2 and Ist1. F. Cartoon of the model for Vps4-triggered sequential disassembly of Vps60-based ESCRT-III polymer.

Importantly, Vps60, unlike Snf7, remained bound to the membrane upon incubation with Vps4/ATP or Ist1-Vps4/ATP (Fig. 5B). Moreover, neither supplementation with Vps4 cofactor Vta1 nor Bro1, which were both suggested to interact with Vps60^74,75^, resulted in a disassembly of Vp60 from the membrane (Fig. 5B). Likewise, Vps60-polymers remained stable with a 10-fold increased Vps4 concentration, which may overcome a lower sensitivity of Vps60-filaments towards Vps4. No direct disassembly of Vps60 by Vps4 in the absence of any downstream ESCRT-III subunits was observed (data not shown).

While Vta1 did not promote Vps4-mediated Vps60 disassembly, it however affected the depolymerization of downstream subunits from Vps60-based polymers. Indeed, Vta1 shifted Ist1’s equilibrium from binding to disassembly (Fig. S5D, E), therefore perturbing Ist1-mediated protection of Did2 from Vps4-mediated disassembly (Fig. S5C, E). Additionally, Vps2 and Vps24 depolymerization rates increased (Fig. 5C-D). Intriguingly, in presence of Vta1, Vps2 and Vps24 depolymerization (Fig. 5 C-D) increased a lot more than Did2’s disassembly did (Fig. 5E, S5C). Overall, these results suggest Vta1 potentially targets specific subunits to increase disassembly. Alternatively, Vta1-binding may primarily increase Vps4-activity, while differences between subunit disassembly rates may rely more on their accessibility to Vps4 within the polymer structure. While all three subunits display simultaneous and synchronic membrane recruitment, Vta1’s strong influence on Vps2 and Vps24 depolymerization rates compared to Did2 establishes a depolymerization hierarchy between these three subunits. It is tempting to speculate that such Vta1-induced divergence of disassembly rates could promote the assembly of ESCRT-III subunits in a temporal sequence (Fig. 5F) as we have previously shown for Snf7-based polymers^41^.

### Vps60- and Snf7-based polymers exert varying and distinct dynamic properties

Vps60 like Snf7 can recruit downstream subunits to form a heteropolymer. To test whether Snf7- and Vps60-polmyers compete during subunit recruitment, we incubated Alexa488-Snf7- and Atto565-Vps60-covered SLBs with Atto647-Vps2, Vps24 and Did2 (Fig. 6A). As a control, we incubated analogous SLBs with Vps2-Vps24, since it binds only to Snf7-patches and not to Vps60-covered membranes (Fig. 6B). Upon recruitment of Vps2-Did2-Vps24, Vps2 fluorescence initially colocalized with Snf7-patches followed by a slower binding to the Vps60-covered membrane (Fig.6A, C, D). In contrast, Vps2-Vps24 only colocalized with Snf7-patches but no recruitment to Vps60-covered membrane was observed (Fig. 6B-D). The two-stepped binding of Vps2-Did2-Vps24, likely emerges from an initial recruitment of Vps2-Vps24 to Snf7-patches. Thereafter, Vps2-Did2-Vp24 bind to Vps60-filaments by an independent recruitment process following slower kinetics (Fig. 3G), implying that recruitment to Snf7- or Vps60-filament occurs independently from each other. To similarly compare disassembly from Snf7 and Vps60 based ESCRT-III polymers, we monitored fluorescence of Vps2 upon addition of Vta1 and Vps4/ATP to SLBs pre-incubated with labeled-Vps2, Vps24, Did2 and Vps60 or Snf7, respectively (Fig. 6E). Vps2 depolymerization from Vps60-based polymers was slightly delayed compared to disassembly from Snf7-based filaments (Fig. 6E).

**Figure 6.**
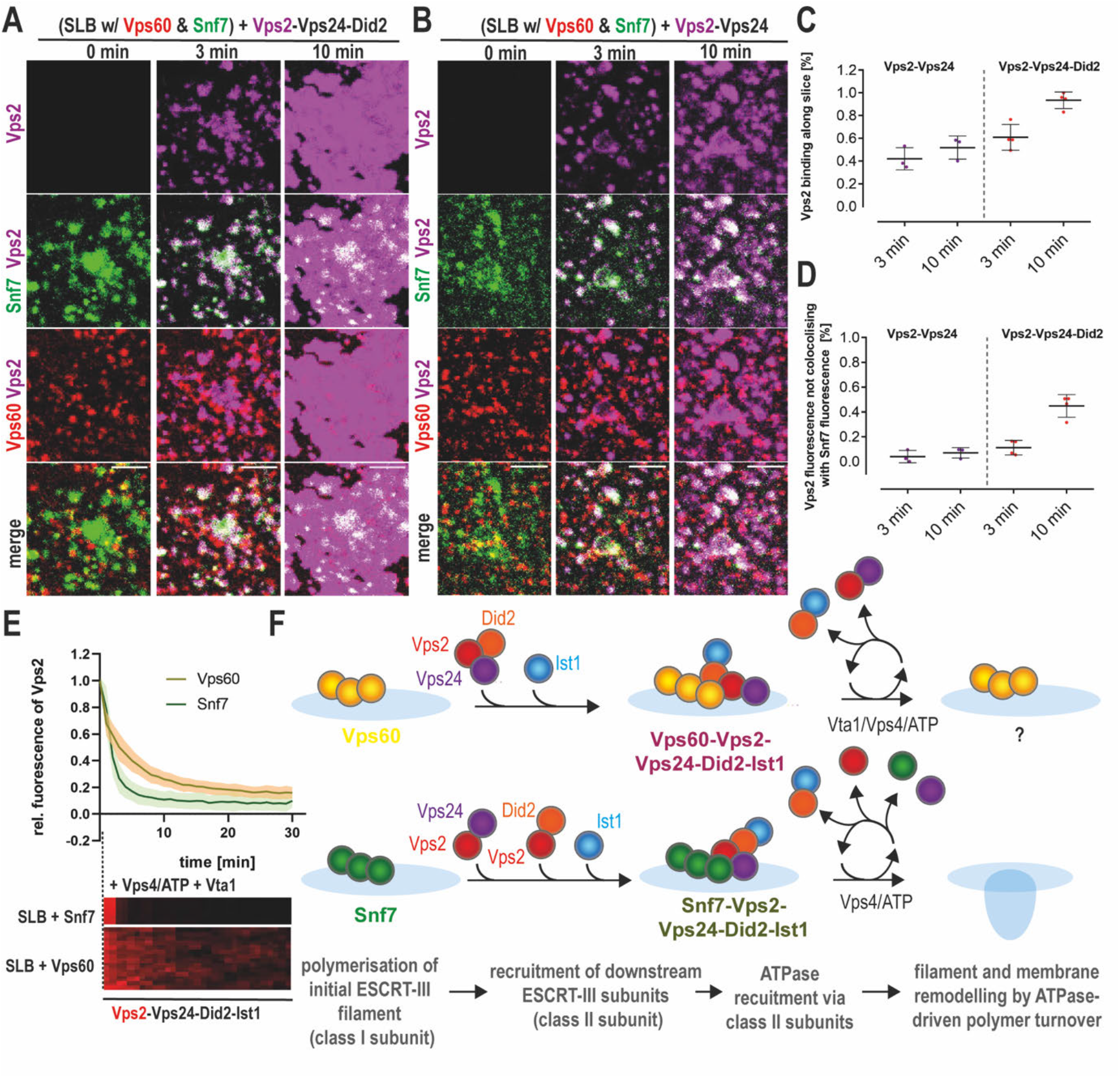
Comparison of Snf7-based and Vps60-based ESCRT-III polymers. A. Confocal images of timelapse experiments of addition of Atto647-Vps2, Vps24 and Did2 to SLBs pre-incubated with Snf7 (green) and Vps60 (red). B. Confocal images of timelapse experiments of addition of Atto647-Vps2 and Vps24 to SLBs pre-incubated with Snf7 (green) and Vps60 (red). C-D Quantification of Vps2 fluorescence intensities total membrane coverage and non-colocalization with SNf7-patches from experiments described in A and B. E. Quantification of Vps2 intensity and kymographs of timelapse experiments of addition of Vta1 and Vps4/ATP to SLBs pre-incubated with Vps2, Vps24, Did1 Ist1 and Snf7 or Vps60. F. Cartoon of proposed model for function of Vps60-based ESCRT-III filaments.

In conclusion, Vps60- and Snf7-based polymers assemble and undergo disassembly independently of each other. These results might indicate that both polymers can co-exist in cells in separated functions. In direct comparison, Vps60-based filaments display delayed assembly and disassembly kinetics which might indicate adaptation to cellular functions in which slower ESCRT-III assembly is required. Alternatively, we might miss cofactors in our in vitro reconstitution approach which could speed up the assembly and disassembly of Vps60-based polymers.

## Discussion

We here show that the ESCRT-III subunit Vps60 functions as the basis for a novel multi-subunit ESCRT-III filament. We further propose this Vps60-based filament to potentially constitute the initiator of a second ESCRT-III polymerization sequence, alternative to the Snf7-based sequence we recently unraveled (Fig. 6F)^41^. In detail, we found Vps60 to polymerize into ring-shaped or curled filaments on membranes, which, in analogy to the Snf7-based polymerization sequence, then recruited ESCRT-III subunits Vps2, Vps24, Did2 and Ist1, before subsequently undergoing Vps4/Vta1-mediated filament turnover. Altogether, our results imply that Vps60, by acting as a template for initiating an alternative ESCRT-III filament, could functionally “replace” Snf7 in specific ESCRT-III functions which require biochemical properties only Vps60-initiated polymers provide^62,65,79,81^.

In support of this notion, a recent study of ESCRT-III in *Plasmodium falciparum* infected red blood cells suggests that PfVps32 (Snf7 homologue) and PfVps60 function in two parallel pathways during formation of extracellular vesicles (EV)^82^. In contradiction to this idea, a very recent study in yeast suggests Vps60 to act downstream of Snf7, Vps2 and Vps24, as recruitment of Vps60 to endosomal membrane was distributed in Snf7, Vps2 and Vps24 knockout mutants^83^. We here, however, find Snf7- and Vps60-polymers to co-exist independently on the same membrane *in vitro*. Additionally, in our assays, in coherence with Banjade *et al*.^66^, Vps60 polymerized spontaneously upon contact with membranes with a higher nucleation rate than Snf7 (this study). Spontaneous nucleation of all established downstream ESCRT-III subunit (Vps2, Vps24, Did2 and Ist1) was absent in our assay.

In contrast to Vps2-Vps24 ^62,68^ which forms the minimal binding unit of Snf7 filaments, recruitment to Vps60 polymers was mediated by Vps2-Did2. This observation highlights the central and pivotal role of Vps2-Did2 during ESCRT-III activity, as already found in our previous study^41^. Overall, our observation of Vps2-Vps24-Did2 recruitment to Vps60 filament is strongly supported by a very recent analysis of ESCRT-III complexes isolated from yeast which reports interactions of Vps60 (Mos10) with Did2, Vps2 and Vps24^83^. Vps4-triggered filament remodeling has previously been established as one of the key factors of ESCRT-III activity in cells^68,79,80^. In our study, Vps60-based filaments were able to recruit Vps4 and underwent partial disassembly. Upon addition of the Vps4 activator Vta1, a disassembly hierarchy between the various subunits was established similarly to sequential depolymerization observed in Snf7-based filament^41,68^. In detail, Vps24 and Vps2 were depolymerized first, followed by Did2 and Ist1. This delay in Did2 disassembly could potentially be mediated by its direct interaction with Vta1^58,59,74–77^. Alternatively, by globally stimulating Vps4 activity, Vta1 might emphasize pre-existing biases in subunit susceptibility to Vps4 e.g., their accessibility within filaments. In stark contrast to Snf7-based polymers, Vps60 polymers are not disassembled in any of the conditions we tested. In cells, Vp60-based filaments might be targeted by another MIT-domain containing AAA-ATPase besides Vps4, or our in vitro reconstitution system might lack a crucial cofactor mediating interaction between Vps4 and Vps60.

Overall, the step-wise subunit recruitment and sequential depolymerization of Vps60-based polymers bares clear resemblance to the Snf7-based polymerization sequence we previously described^41^, and suggest the Vps60-based filament described in this study might form the initiation of an alternative Vps60-based ESCRT-III polymerization sequence (Fig. 6 F). Bioinformatic analysis of ESCRT-III proteins classes the subunits into two groups according to their domain conservation: the Snf7 family, Vps20, Snf7 and Vps60, groups clearly the nucleators and initiators of ESCRT-III (class I), and the Vps2 family encompassing Vps2, Vps24 and Did2 (class II), groups the “pivots” which recruit Vps4^57^. At a closer look, both Vps60-based and Snf7-based ESCRT-III polymers and their corresponding presumed polymerization sequences share remarkable similarities regarding their organization, indicating that ESCRT-III-mediated membrane remodeling might follow a general mechanism (Fig. 6F). An initial class I polymer (ring or spiral) mediates binding of class II subunits (helices) which then recruit an ATPase to trigger filament remodeling to promote ESCRT-III activity. In fact, this generalized progression of events is supported by the observation that direct recruitment of Vps4 via an initiating class I protein is non-functional *in vivo*^66^, potentially due to premature disassembly of the ESCRT-III polymer^66^ or a lack of filament remodeling capacity^55^.

A remaining, yet essential question about the Vps60-based dynamic ESCRT-III polymers is their cellular function. Overall, the specific differences such as structure and kinetics between both ESCRT-III polymers and their associated sequences hint towards functional specialization of the filaments, adopting requirements of distinct cellular functions. This notion, is supported by a unique cellular localization of GFP-tagged Vps60 *in vivo* to not only the endosomal, as it is observed for Snf7, but also yeast’s vacuolar membrane^66,83^. Separated functional pathways were also suggested for ESCRT-III subunits PfVps32 (Snf7 homolog) and PfVps60 during EV-formation in *Plasmodium falciparum* infected red blood cells^82^. Finally, the author attributed the formation of smaller vesicles to the Vps60-dependent pathway which would agree with the smaller filament structures we observed.

## Acknowledgements

A.R. acknowledges funding from the Swiss National Fund for Research Grants N°31003A_130520, N°31003A_149975 and N°31003A_173087. The authors want to thank the NCCR Chemical Biology for constant support during this project.

## Authors contributions

A.R. and A.-K.P. conceptualized the study. A.-K.P. and H.Z. designed, performed and analyzed experiments. H.Z. and F.H. purified proteins. A.-K.P. and A.R. wrote the manuscript with help from H.Z.

## Methods

### Protein purification

Vps60 (pGEX4T-Vps60) and Vta1 (pGEX4T-Vta1) (gift from David Katzmann lab, Mayo clinic, USA) were expressed for 4h in Bl21 *e*.*coli* following IPTG induction. Bacteria were lysed (1% triton, 20 mM Hepes pH 8.0, 150 mM NaCl, cOmplete) using sonication and soluble lysate was loaded on GST-resin (washing buffer: 20 mM Hepes, pH 8.0, 150 NaCl), before on-column cleavage using TEV-protease was performed. Vta1-GST was eluted prior to TEV treatment as on-column digest was inefficient (elution buffer: 150 mM NaCl, 10 mM glutathione, 20 mM Hepes pH 8.0). TEV protease was removed using Ni-NTA resin and the purified protein of interest was dialyzed against storage buffer (20 mM Hepes, pH 8.0) before concentration.

Increase of Vps4 ATPase-activity upon addition of Vta1, was checked by malachite green assay at the end of each purification to confirm the proper functioning of Vta1. In short, a malachite green stock solution was prepared by mixing 3 parts of 0.045 % malachite green in water with 2 parts of 4.2% Ammonium Molybdate in 4 M HCl for 1 hour under constant stirring before supplementation with 0.01 % Tween20.For quantification a calorimetric measurement at 620nm was performed 5min after mixing of 3 parts malachite green stock solution with 1 part of reaction solution either resulting from incubation of Vps4 and Vta1in reaction buffer (20 mM Hepes, 150 mM NaCl, 2mM MgCl2 pH=7.5) supplemented with 1mM ATP or a phosphate calibration curve.

Following the labelling procedure given by the reagent provider, Snf7, Vps2, Ist1 and Vps24 were labeled with TFP-AlexaFluor-488 (Ref N°A-30005, ThermoFisher Scientific,). Vps2 and Vps60 were labeled with TFP-Atto-565 (Atto-Tec AD 565-3). Did2 was labeled with maleimide-AlexFluor-488 (ThermoFisher Scientific, A-30005). Vps2 was labeled with NHS-Atto-647N (Atto-Tec AD 647N). If not otherwise mentioned, following protein concentration were used: ESCRT-II 1μM, Vps20 1μM, Bro1 500nM, Vps60 50nM, Snf7 400nM, Vps2 1 μM, Vps24 1 μM, Did2 1 μM, Ist1 1 μM, Vps4 1 μM, Vta1 500nM ATP 2 mM. In general, labeled proteins were mixed 1:1 with unlabeled protein.

### Preparation of giant unilamellar vesicles (GUV) and large giant unilamellar vesicles (LUV)

GUVs were prepared by electroformation: 20-30 μL of a 2mg/ml lipid solution in chloroform (DOPC:DOPS:DOPE-Atto647N:DSPE-PEG(2000)Biotin, 6:4:0.01:0.003; Avanti Polar Lipids, Atto-tec) were dried on indium-tin oxide (ITO)-coated glass slides for 1h. For experiments including Atto647N-Vps2 unlabeled GUVs were used. A growth chamber was assembled by clamping a rubber ring between the ITO-slides, filled with 500 μl of a sucrose buffer osmotically equilibrated with the experimental buffer. ITO-Slides were then connected to an AC generator set under 1V AC (10 Hz) for 1.5h. GUVs were stored at 4°C for at maximum a week.

For LUV preparation, DOPC:DOPS (6:4; 10 mg/ml) mixture was evaporated in a glass tube, 500 μl of buffer were added, the tube was vortex followed by 5 times freezing and thawing. LUVs were stored at -20 °C and extruded with a 200 nm filter before usage.

### Supported membrane bilayer assay

Supported membrane bilayer assay was performed as described in ^46^. Experiments were performed in 20 mM Tris pH.6.8, 200 mM NaCl and 1mM MgCl_2_. 2 mM DTT was added to the buffer for experiments including Ist1. GUVs diluted in buffer were burst on a plasma-cleaned coverslip forming the bottom of a flow chamber (coverslip and sticky-Slide VI 0.4, Ibidi) to form supported bilayers. Thereafter, the chamber was passivated with Casein (1mg/ml Sigma -Aldrich) for 10 min and washed with buffer, before the experiments was conducted. Subsequent changes of protein or buffer solutions in the chamber were made via a syringe pump connected to the flow chamber. Briefly, protein preparations are diluted in reaction buffer with 80 μL final volume. In presence of Ist and Did2, 0.8 μL DTT 1 mM is added. And reaction buffer has 2 mM MgCl_2_ added. All proteins have a final concentration of 1 μM during acquisitions except Vps60 and Snf7. Snf7 patches are pre-grown at 400 nM until, then Snf7 is washed out before other protein is added. Vps60 was tested with many concentrations for dynamics assays and added at 50 nM for 9 minutes when pre-grown for experiments.

Partially adhered vesicles were prepared as described in ^46^. Briefly, a flow chamber assembled from a coverslip and sticky-Slide VI 0.4, Ibidi was incubated with Avidin (0.1 mg/ml) for 10 min, before washing with buffer (20 mM Tris pH.6.8, 200 mM NaCl and 1mM MgCl_2_) and addition of GUVs (including 0.03 % DSPE-PEG(2000)Biotin) diluted in buffer. As soon as GUVs started to attach biotinylated-Albumin (1mg/ml, Sigma-Aldrich) was added to stop attachment and prevent bursting of the GUVs.

### Image acquisition

Confocal Imaging was performed on an inverted spinning disc microscope assembled by 3i (Intelligent Imaging Innovation) consisting of a Nikon base (Eclipse C1, Nikon), a 100x 1.49 NA oil immersion objective and an EVOLVE EM-CCD camera (Ropper Scientific Inc.). For analysis of supported bilayer experiments, 3 μm thick Z-stack were maximally projected using a Fiji plugin ^84^. X-y drift of the microscopy was corrected using the plugin Turboreg and a custom-written ImageJ macro. For artificial membrane neck experiments, 15 μm thick Z-stacks were acquired.

### Electron microscopy

For EM experiments, LUVs were diluted 1:100 in buffer (20 mM Tris pH.6.8, 200 mM NaCl and 1mM MgCl_2_), spun down (10’, 5,000g), resuspended in 250 nM Vps60 for 1h at 4°C. Samples were absorbed onto Carbon-coated grids Cu 300 and stained with 2% uranyl acetate for 30s. Images were acquired on a Tecnai G2 Sphera (FEI) electron microscope.

### Optical tweezer tube pulling experiment

Membrane nanotube pulling experiments were performed on the setup published in ^46^ allowing simultaneous optical tweezer application, spinning disc confocal and brightfield imaging based on an inverted Nikon eclipse Ti microscope and a 5W 1064nm laser focused through a 100 × 1.3 NA oil objective (ML5-CW-P-TKS-OTS, Manlight). Membrane nanotubes were pulled with streptavidin beads (3.05 μm, Spherotec) from a GUV containing 0.01% DSPE-PEG(2000)Biotin and aspired in a motorized micropipette (MP-285, Sutter Instrument). Proteins were injected using a slightly bigger micropipette connected to a pressure control system (MFCS-VAC -69 mbar, Fluigent).

### Quantification and statistical analysis

For quantification of supported bilayer experiments, integrated fluorescence intensity of membrane patches (Vps60) of single proteins patches (Snf7) was measured using Fiji, background at time 0 min subtracted, normalized to time point 0 and a kymograph was extracted for dynamic experiments. Fluorescence intensities were normalized by their maximum value. To determine the colocalization of Atto565-Vps60 and Alexa488-Snf7 or Atto647-Vps2 and Atto565-Vps60 or Alexa488-Snf7, relative fluorescence was measured along linearized membrane contours, relative fluorescence values were binarized (1 above threshold, 0 below, thresholds: 0.3 Snf7, 0.4 DOPE). The percentage of no colocalization was extracted by the proportion of pixels with the value 1 from the Snf7 channel for which the value in the membrane channel was 0. No colocalization was only counted at a minimal distance of four pixels to the nearest membrane neck (value 1).

For quantification of nucleation rates, membrane areas were isolated from images and Vps60-puncta or total fluorescence of Vps60 after background subtraction (t = 0min) was extracted and dived through total membrane area.

For all experiments the mean and standard deviation (SD) were calculated. Number of independent experiments (n) und number of patches or membrane necks (ROI) analyzed are indicated in the corresponding figure legends. The graph and statistics were done using Prism 8 (GraphPad software).

## Extended Data Figures

**Figure S1.**
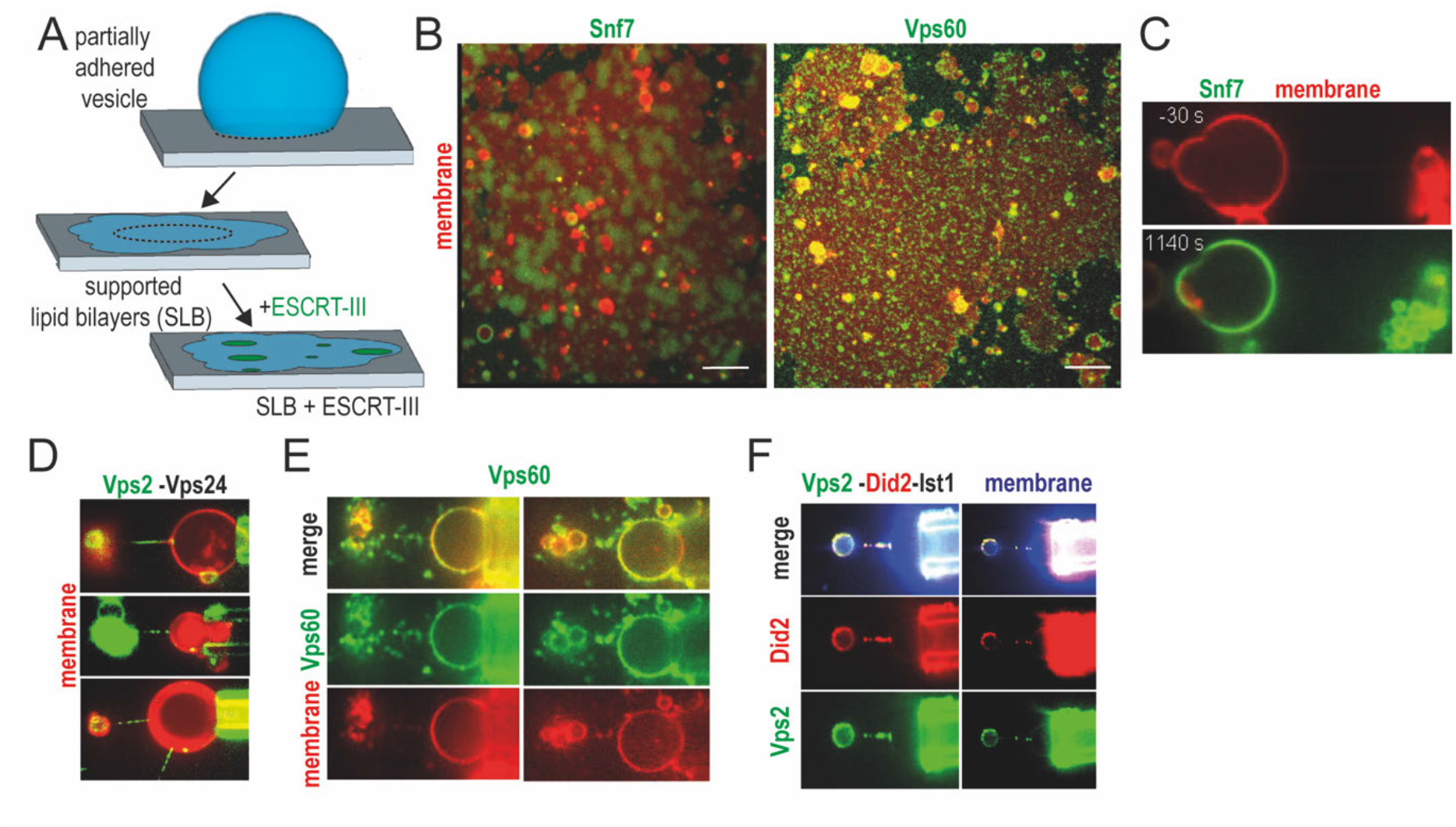
Characterization of ESCRT-III protein properties. A. Schematic presentation of SLBs formation. B. Confocal images of Snf7 (green) o. Vps60 (green) nucleation on SLBs (red). C. Confocal microscopy images of Snf7-Alex488 (green) binding to membrane nanotubes (red). D. Confocal microscopy images of Vps60 (green) binding to membrane nanotubes (red). E. Confocal microscopy images of Vps2-Alex488 (green) and Vps24 binding to membrane nanotubes (red). F. Confocal microscopy images of Vps2-Alex488 (green), Did2-Atto565 (red) and Ist1 binding to membrane nanotubes (blue).

**Figure S2.**
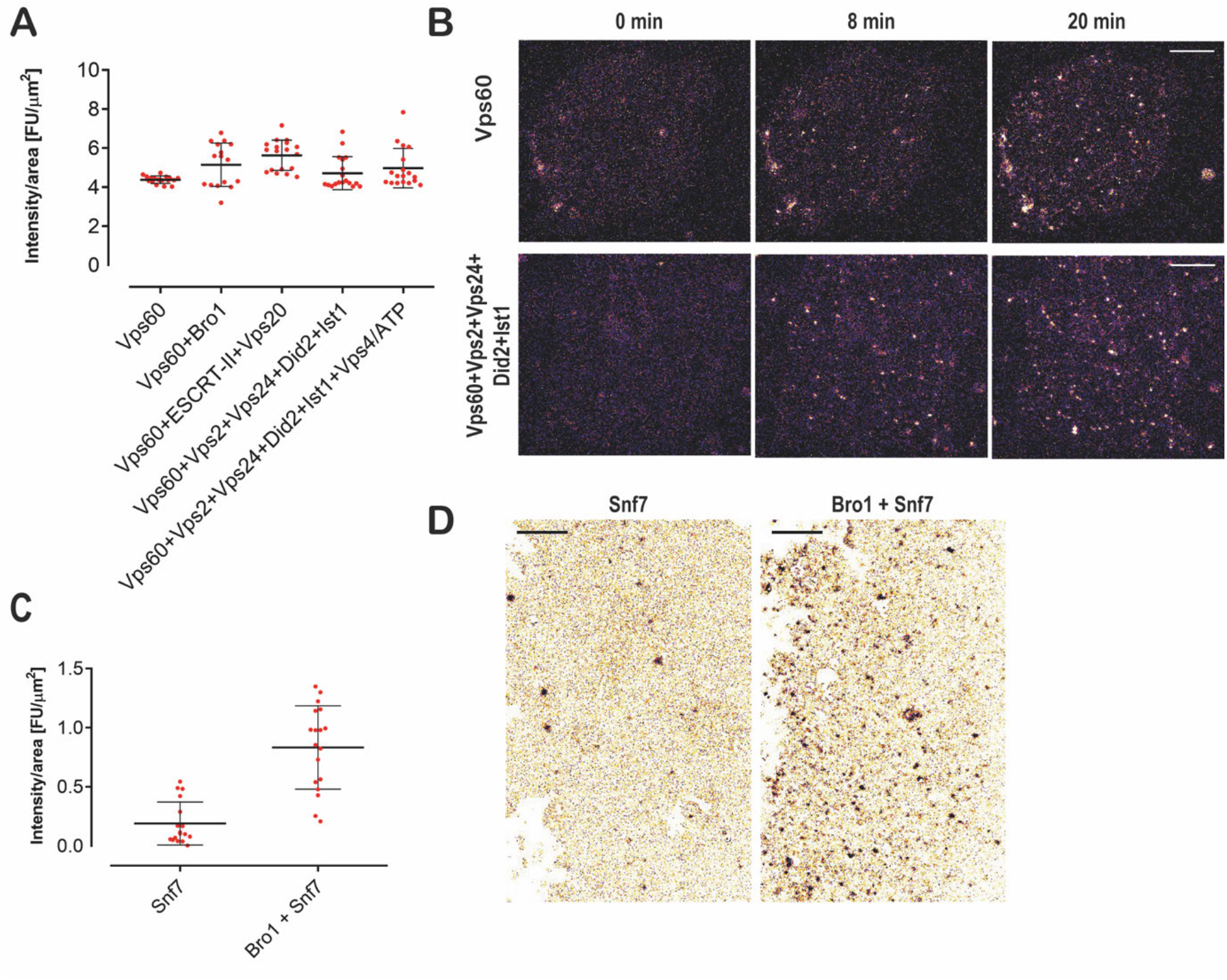
Vps60 nucleation pattern on SLBs. A. Quantification Vps60 intensity from experiments described in Fig. 2A. B. Confocal images of timelapse experiments of SLBs incubated with the indicated mixture of proteins (scale bar 10 μm). C-D. Quantification of Snf7 intensity (C) and confocal images (D) of SLBs incubated with Alexa488-Snf7 or Alexa488-Snf7 and Bro1.

**Figure S3.**
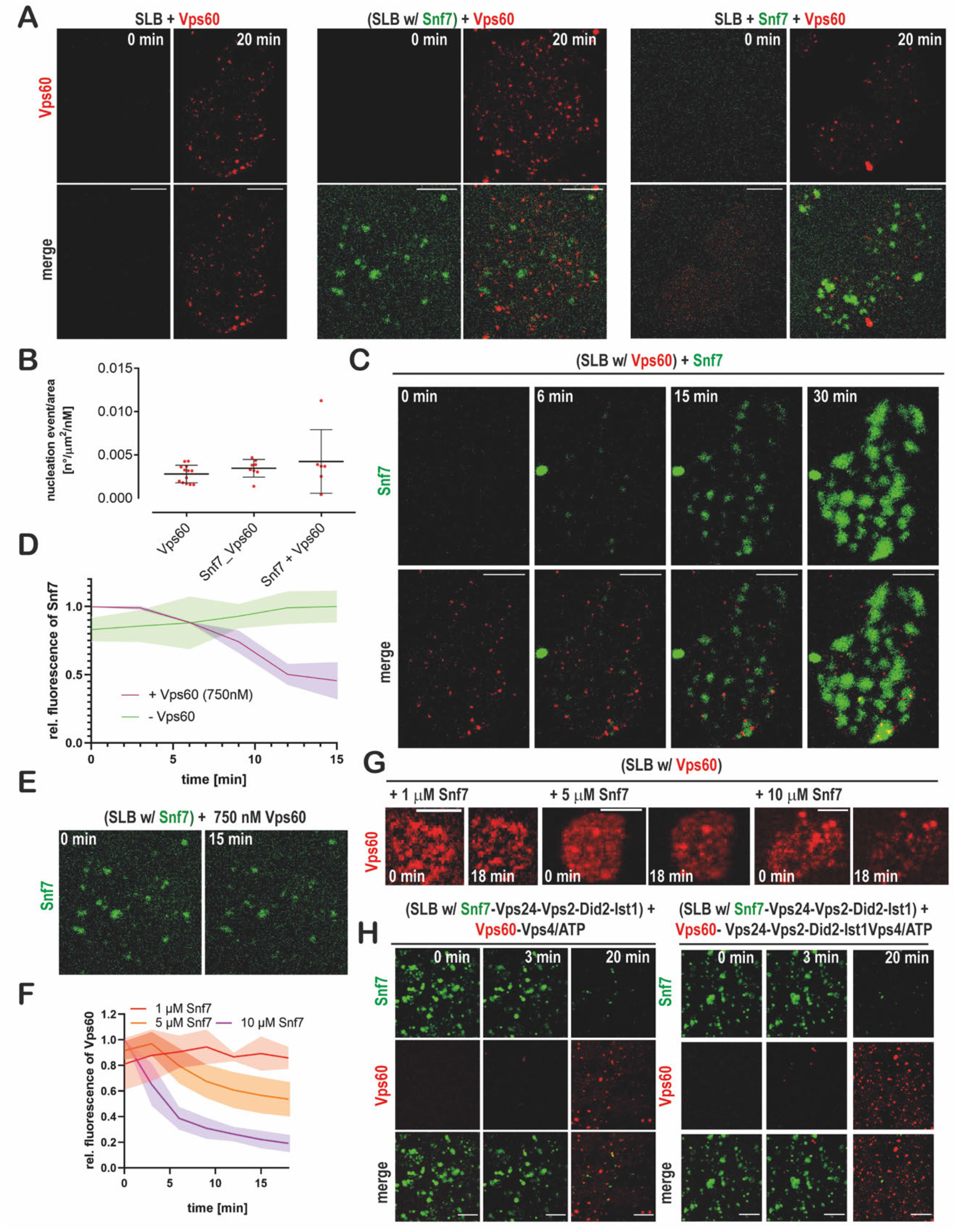
Vps60 and Snf7 bind membrane independently from each other. A. Confocal images from experiments described in Fig. 3A. B. Quantification of Vps60 intensity from experiments described in Fig. 3A. C. Confocal images of timelapse experiments of addition of Alexa488-Snf7 to Vps60-covered SLBs. D-E. Quantification of Snf7 intensity (E) and confocal images (D) of timelapse experiments of addition of Vps60 (unlabelled) to SLBs with pre-grown Snf7-patches.

**Figure S4.**
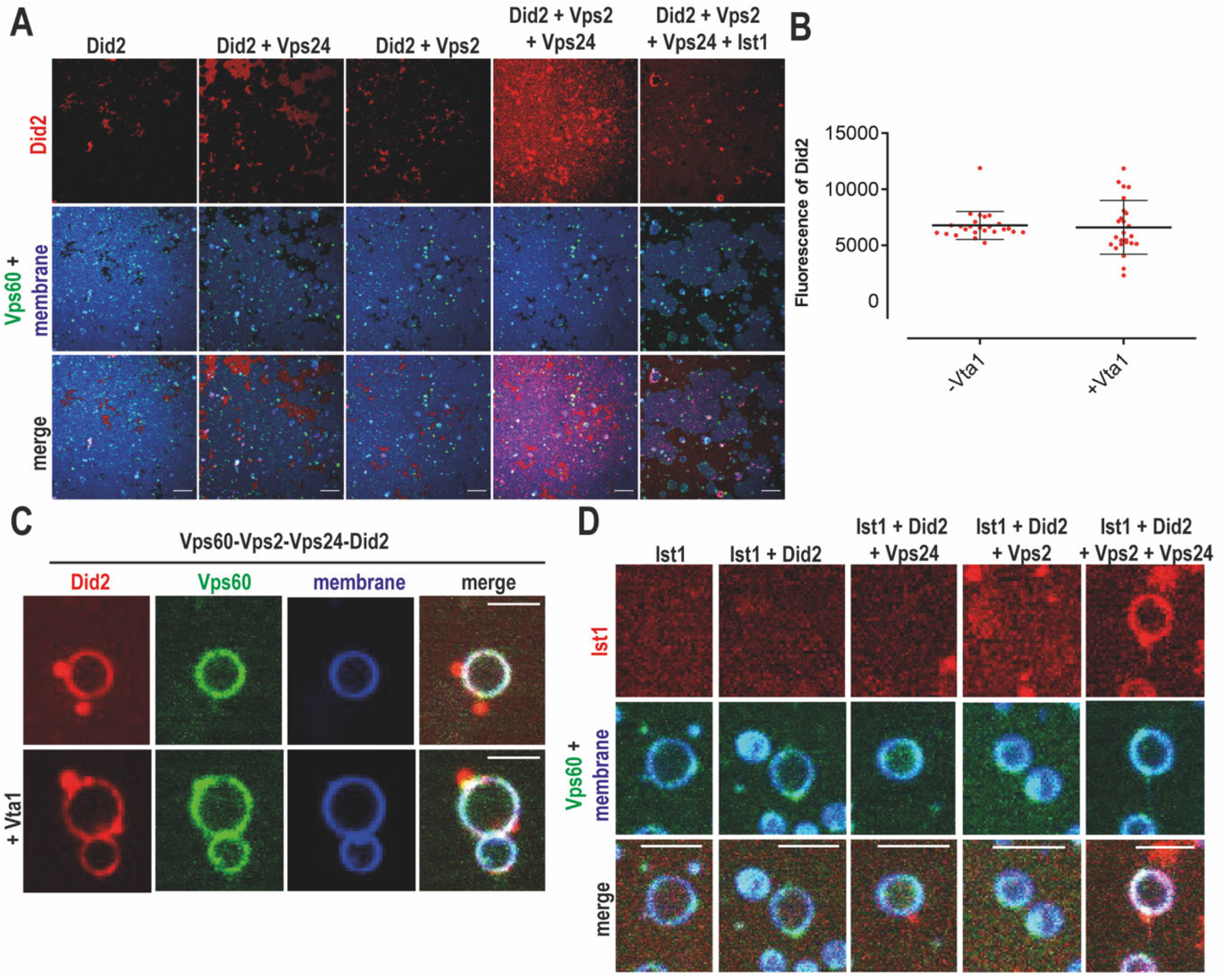
Membrane-bound Vps60 recruits ESCRT-III proteins. A. Confocal images of experiments described in Fig. 4E. B-C. Quantification of Did2 intensity (B) and confocal images (C) of Vps60-coverd SLBs incubated with Vps2, Vps24, Alexa488-Did2 and Ist1 in presence or absence of Vta1. D. Confocal images of experiments described in Fig. 4F.

**Figure S5.**
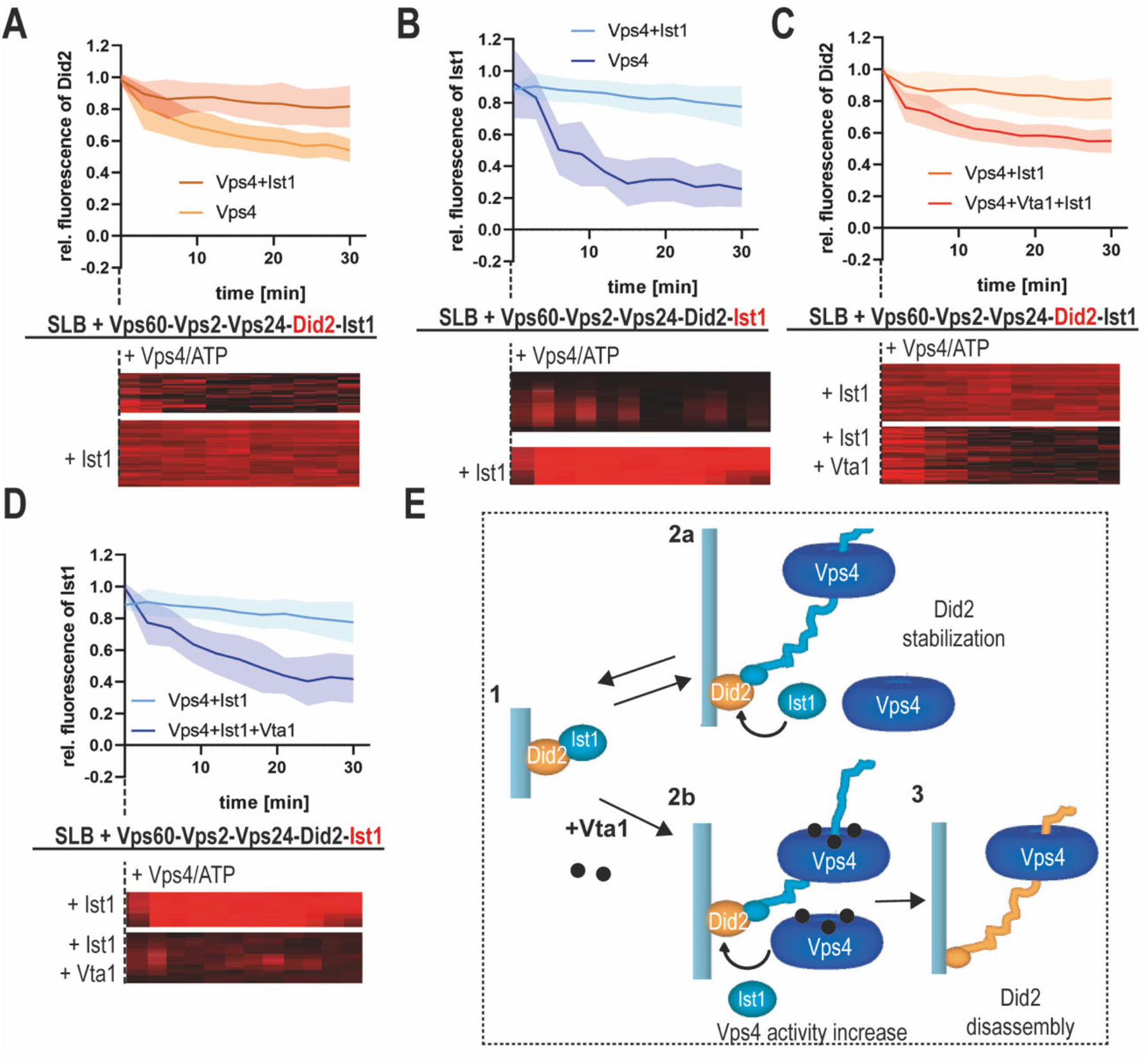
Vps4-triggered depolymerization of Did2 and Ist1 from Vps60-based ESCRT-III filaments. A. Quantification of Did2 intensity and kymographs of timelapse experiments of addition of Vps4/ATP or Ist1 and Vps4/ATP to SLBs pre-incubated with Vps60, Vps2, Vps24, Alexa488-Did2 and Ist1. B. Quantification of Ist1 intensity and kymographs of timelapse experiments of addition of Vps4/ATP or Ist1 and Vps4/ATP to SLBs pre-incubated with Vps60, Vps2, Vps24, Did2 and Alexa488-Ist1. C. Quantification of Did2 intensity and kymographs of timelapse experiments of addition of Ist1 and Vps4/ATP or Ist1, Vps4/ATP or Vta1 to SLBs pre-incubated with Vps60, Vps2, Vps24, Alexa488-Did2 and Ist1. B. Quantification of Ist1 intensity and kymographs of timelapse experiments of addition of Ist1 and Vps4/ATP or Ist1, Vps4/ATP or Vta1 to SLBs pre-incubated with Vps60, Vps2, Vps24, Did2 and Alexa488-Ist1. E. Cartoon of the model for interplay of Vps4, Did2 and Ist1 during disassembly of ESCRT-III polymers in presence or absence of Vta1.

## References

1. Spang, A. et al. Complex archaea that bridge the gap between prokaryotes and eukaryotes. Nature 521, 173–179 (2015).

2. Liu, J. et al. Bacterial Vipp1 and PspA are members of the ancient ESCRT-III membrane-remodeling superfamily. Cell (2021) doi:10.1016/j.cell.2021.05.041.

3. Schöneberg, J., Lee, I.-H., Iwasa, J. H. & Hurley, J. H. Reverse-topology membrane scission by the ESCRT proteins. Nat. Rev. Mol. Cell Biol. 18, 5–17 (2017).

4. Gatta, A. T. & Carlton, J. G. The ESCRT-machinery: closing holes and expanding roles. Curr Opin Cell Biol 59, 121–132 (2019).

5. Vietri, M., Radulovic, M. & Stenmark, H. The many functions of ESCRTs. Nat Rev Mol Cell Biol 21, 25–42 (2020).

6. Barnes, J. & Wilson, D. W. Seeking Closure: How Do Herpesviruses Recruit the Cellular ESCRT Apparatus? J. Virol. 93, (2019).

7. Broniarczyk, J. et al. The VPS4 component of the ESCRT machinery plays an essential role in HPV infectious entry and capsid disassembly. Sci Rep 7, 45159 (2017).

8. Lippincott-Schwartz, J., Freed, E. O. & van Engelenburg, S. B. A Consensus View of ESCRT-Mediated Human Immunodeficiency Virus Type 1 Abscission. Annu Rev Virol 4, 309–325 (2017).

9. Ortmann, A. C. et al. Transcriptome analysis of infection of the archaeon Sulfolobus solfataricus with Sulfolobus turreted icosahedral virus. J. Virol. 82, 4874–4883 (2008).

10. Streck, N. T., Carmichael, J. & Buchkovich, N. J. Nonenvelopment Role for the ESCRT-III Complex during Human Cytomegalovirus Infection. J. Virol. 92, (2018).

11. Tabata, K. et al. Unique Requirement for ESCRT Factors in Flavivirus Particle Formation on the Endoplasmic Reticulum. Cell Rep 16, 2339–2347 (2016).

12. Denais, C. M. et al. Nuclear envelope rupture and repair during cancer cell migration. Science 352, 353–358 (2016).

13. Jimenez, A. J. et al. ESCRT machinery is required for plasma membrane repair. Science 343, 1247136 (2014).

14. Raab, M. et al. ESCRT III repairs nuclear envelope ruptures during cell migration to limit DNA damage and cell death. Science 352, 359–362 (2016).

15. Radulovic, M. et al. ESCRT-mediated lysosome repair precedes lysophagy and promotes cell survival. EMBO J. 37, (2018).

16. Skowyra, M. L., Schlesinger, P. H., Naismith, T. V. & Hanson, P. I. Triggered recruitment of ESCRT machinery promotes endolysosomal repair. Science 360, (2018).

17. Junglas, B. et al. PspA adopts an ESCRT-III-like fold and remodels bacterial membranes. Cell (2021) doi:10.1016/j.cell.2021.05.042.

18. Gupta, T. K. et al. Structural basis for VIPP1 oligomerization and maintenance of thylakoid membrane integrity. Cell (2021) doi:10.1016/j.cell.2021.05.011.

19. Göser, V., Kehl, A., Röder, J. & Hensel, M. Role of the ESCRT-III complex in controlling integrity of the Salmonella-containing vacuole. Cell. Microbiol. e13176 (2020) doi:10.1111/cmi.13176.

20. López-Jiménez, A. T. et al. The ESCRT and autophagy machineries cooperate to repair ESX-1-dependent damage at the Mycobacterium-containing vacuole but have opposite impact on containing the infection. PLoS Pathog. 14, e1007501 (2018).

21. Allison, R. et al. An ESCRT-spastin interaction promotes fission of recycling tubules from the endosome. J. Cell Biol. 202, 527–543 (2013).

22. Chang, C.-L. et al. Spastin tethers lipid droplets to peroxisomes and directs fatty acid trafficking through ESCRT-III. J. Cell Biol. (2019) doi:10.1083/jcb.201902061.

23. Mast, F. D. et al. ESCRT-III is required for scissioning new peroxisomes from the endoplasmic reticulum. J. Cell Biol. 217, 2087–2102 (2018).

24. Dores, M. R., Grimsey, N. J., Mendez, F. & Trejo, J. ALIX Regulates the Ubiquitin-Independent Lysosomal Sorting of the P2Y1 Purinergic Receptor via a YPX3L Motif. PLoS One 11, e0157587 (2016).

25. Dores, M. R. et al. AP-3 regulates PAR1 ubiquitin-independent MVB/lysosomal sorting via an ALIX-mediated pathway. Mol Biol Cell 23, 3612–3623 (2012).

26. Pashkova, N. et al. The yeast Alix homolog Bro1 functions as a ubiquitin receptor for protein sorting into multivesicular endosomes. Dev Cell 25, 520–533 (2013).

27. Olmos, Y., Perdrix-Rosell, A. & Carlton, J. G. Membrane Binding by CHMP7 Coordinates ESCRT-III-Dependent Nuclear Envelope Reformation. Curr Biol 26, 2635–2641 (2016).

28. Vietri, M. et al. Spastin and ESCRT-III coordinate mitotic spindle disassembly and nuclear envelope sealing. Nature 522, 231–235 (2015).

29. Webster, B. M. et al. Chm7 and Heh1 collaborate to link nuclear pore complex quality control with nuclear envelope sealing. EMBO J 35, 2447–2467 (2016).

30. Babst, M., Katzmann, D. J., Estepa-Sabal, E. J., Meerloo, T. & Emr, S. D. Escrt-III: an endosome-associated heterooligomeric protein complex required for mvb sorting. Dev. Cell 3, 271–282 (2002).

31. Teis, D., Saksena, S. & Emr, S. D. Ordered assembly of the ESCRT-III complex on endosomes is required to sequester cargo during MVB formation. Dev. Cell 15, 578–589 (2008).

32. Teis, D., Saksena, S., Judson, B. L. & Emr, S. D. ESCRT-II coordinates the assembly of ESCRT-III filaments for cargo sorting and multivesicular body vesicle formation. EMBO J. 29, 871–883 (2010).

33. A Saksena, S., Wahlman, J., Teis, D., Johnson, A. E. & Emr, S. D. Functional reconstitution of ESCRT-III assembly and disassembly. Cell 136, 97–109 (2009).

34. Obita, T. et al. Structural basis for selective recognition of ESCRT-III by the AAA ATPase Vps4. Nature 449, 735–739 (2007).

35. Stuchell-Brereton, M. D. et al. ESCRT-III recognition by VPS4 ATPases. Nature 449, 740–744 (2007).

36. Brune, T., Kunze-Schumacher, H. & Kölling, R. Interactions in the ESCRT-III network of the yeast Saccharomyces cerevisiae. Curr Genet 65, 607–619 (2019).

37. Lata, S. et al. Helical structures of ESCRT-III are disassembled by VPS4. Science 321, 1354–1357 (2008).

38. Adell, M. A. Y. et al. Coordinated binding of Vps4 to ESCRT-III drives membrane neck constriction during MVB vesicle formation. J. Cell Biol. 205, 33–49 (2014).

39. Yang, B., Stjepanovic, G., Shen, Q., Martin, A. & Hurley, J. H. Vps4 disassembles an ESCRT-III filament by global unfolding and processive translocation. Nat. Struct. Mol. Biol. 22, 492–498 (2015).

40. Mierzwa, B. E. et al. Dynamic subunit turnover in ESCRT-III assemblies is regulated by Vps4 to mediate membrane remodelling during cytokinesis. Nat. Cell Biol. 19, 787–798 (2017).

41. Pfitzner, A.-K. et al. An ESCRT-III Polymerization Sequence Drives Membrane Deformation and Fission. Cell 182, 1140–1155.e18 (2020).

42. Guizetti, J. et al. Cortical constriction during abscission involves helices of ESCRT-III-dependent filaments. Science 331, 1616–1620 (2011).

43. Adell, M. A. Y. et al. Recruitment dynamics of ESCRT-III and Vps4 to endosomes and implications for reverse membrane budding. Elife 6, (2017).

44. A Henne, W. M., Buchkovich, N. J., Zhao, Y. & Emr, S. D. The endosomal sorting complex ESCRT-II mediates the assembly and architecture of ESCRT-III helices. Cell 151, 356–371 (2012).

45. Shen, Q.-T. et al. Structural analysis and modeling reveals new mechanisms governing ESCRT-III spiral filament assembly. J. Cell Biol. 206, 763–777 (2014).

46. Chiaruttini, N. et al. Relaxation of Loaded ESCRT-III Spiral Springs Drives Membrane Deformation. Cell 163, 866–879 (2015).

47. Hanson, P. I., Roth, R., Lin, Y. & Heuser, J. E. Plasma membrane deformation by circular arrays of ESCRT-III protein filaments. J. Cell Biol. 180, 389–402 (2008).

48. Effantin, G. et al. ESCRT-III CHMP2A and CHMP3 form variable helical polymers in vitro and act synergistically during HIV-1 budding. Cell. Microbiol. 15, 213–226 (2013).

49. Nguyen, H. C. et al. Membrane constriction and thinning by sequential ESCRT-III polymerization. Nat Struct Mol Biol 27, 392–399 (2020).

50. McCullough, J. et al. Structure and membrane remodeling activity of ESCRT-III helical polymers. Science 350, 1548–1551 (2015).

51. Hanson, P. I., Roth, R., Lin, Y. & Heuser, J. E. Plasma membrane deformation by circular arrays of ESCRT-III protein filaments. J Cell Biol 180, 389–402 (2008).

52. Moser von Filseck, J. et al. Anisotropic ESCRT-III architecture governs helical membrane tube formation. Nat Commun 11, 1516 (2020).

53. Bertin, A. et al. Human ESCRT-III polymers assemble on positively curved membranes and induce helical membrane tube formation. Nat Commun 11, 2663 (2020).

54. Harker-Kirschneck, L. et al. Physical mechanisms of ESCRT-III-driven cell division. Proc Natl Acad Sci U S A 119, e2107763119 (2022).

55. Pfitzner, A.-K., Moser von Filseck, J. & Roux, A. Principles of membrane remodeling by dynamic ESCRT-III polymers. Trends in Cell Biology (2021) doi:10.1016/j.tcb.2021.04.005.

56. Katzmann, D. J., Babst, M. & Emr, S. D. Ubiquitin-dependent sorting into the multivesicular body pathway requires the function of a conserved endosomal protein sorting complex, ESCRT-I. Cell 106, 145–155 (2001).

57. Leung, K. F., Dacks, J. B. & Field, M. C. Evolution of the multivesicular body ESCRT machinery; retention across the eukaryotic lineage. Traffic 9, 1698–1716 (2008).

58. Nickerson, D. P., West, M., Henry, R. & Odorizzi, G. Regulators of Vps4 ATPase activity at endosomes differentially influence the size and rate of formation of intralumenal vesicles. Mol Biol Cell 21, 1023–1032 (2010).

59. Azmi, I. F. et al. ESCRT-III family members stimulate Vps4 ATPase activity directly or via Vta1. Dev Cell 14, 50–61 (2008).

60. Chiaruttini, N. et al. Relaxation of Loaded ESCRT-III Spiral Springs Drives Memsbrane Deformation. Cell 163, 866–879 (2015).

61. De Franceschi, N. et al. The ESCRT protein CHMP2B acts as a diffusion barrier on reconstituted membrane necks. J Cell Sci 132, (2018).

62. Teis, D., Saksena, S. & Emr, S. D. Ordered assembly of the ESCRT-III complex on endosomes is required to sequester cargo during MVB formation. Dev Cell 15, 578–589 (2008).

63. Lata, S. et al. Structural basis for autoinhibition of ESCRT-III CHMP3. J Mol Biol 378, 818–827 (2008).

64. Effantin, G. et al. ESCRT-III CHMP2A and CHMP3 form variable helical polymers in vitro and act synergistically during HIV-1 budding. Cell Microbiol 15, 213–226 (2013).

65. Henne, W. M., Buchkovich, N. J., Zhao, Y. & Emr, S. D. The endosomal sorting complex ESCRT-II mediates the assembly and architecture of ESCRT-III helices. Cell 151, 356–371 (2012).

66. Banjade, S., Shah, Y. H., Tang, S. & Emr, S. D. Design principles of the ESCRT-III Vps24-Vps2 module. Elife 10, (2021).

67. Lenz, M., Crow, D. J. G. & Joanny, J.-F. Membrane buckling induced by curved filaments. Phys Rev Lett 103, 038101 (2009).

68. Mierzwa, B. E. et al. Dynamic subunit turnover in ESCRT-III assemblies is regulated by Vps4 to mediate membrane remodelling during cytokinesis. Nat Cell Biol 19, 787–798 (2017).

69. Tang, S. et al. ESCRT-III activation by parallel action of ESCRT-I/II and ESCRT-0/Bro1 during MVB biogenesis. Elife 5, (2016).

70. Fyfe, I., Schuh, A. L., Edwardson, J. M. & Audhya, A. Association of the endosomal sorting complex ESCRT-II with the Vps20 subunit of ESCRT-III generates a curvature-sensitive complex capable of nucleating ESCRT-III filaments. J Biol Chem 286, 34262–34270 (2011).

71. Larios, J., Mercier, V., Roux, A. & Gruenberg, J. ALIX-and ESCRT-III-dependent sorting of tetraspanins to exosomes. J Cell Biol 219, (2020).

72. Teis, D., Saksena, S., Judson, B. L. & Emr, S. D. ESCRT-II coordinates the assembly of ESCRT-III filaments for cargo sorting and multivesicular body vesicle formation. EMBO J 29, 871–883 (2010).

73. Mu, R. et al. Two distinct binding modes define the interaction of Brox with the C-terminal tails of CHMP5 and CHMP4B. Structure 20, 887–898 (2012).

74. Yang, Z. et al. Structural basis of molecular recognition between ESCRT-III-like protein Vps60 and AAA-ATPase regulator Vta1 in the multivesicular body pathway. J Biol Chem 287, 43899–43908 (2012).

75. Shen, J. et al. NMR studies on the interactions between yeast Vta1 and Did2 during the multivesicular bodies sorting pathway. Sci Rep 6, 38710 (2016).

76. Monroe, N., Han, H., Shen, P. S., Sundquist, W. I. & Hill, C. P. Structural basis of protein translocation by the Vps4-Vta1 AAA ATPase. Elife 6, (2017).

77. Norgan, A. P. et al. Relief of autoinhibition enhances Vta1 activation of Vps4 via the Vps4 stimulatory element. J Biol Chem 288, 26147–26156 (2013).

78. Rue, S. M., Mattei, S., Saksena, S. & Emr, S. D. Novel Ist1-Did2 complex functions at a late step in multivesicular body sorting. Mol Biol Cell 19, 475–484 (2008).

79. Adell, M. A. Y. et al. Recruitment dynamics of ESCRT-III and Vps4 to endosomes and implications for reverse membrane budding. Elife 6, (2017).

80. Guizetti, J. et al. Cortical constriction during abscission involves helices of ESCRT-III-dependent filaments. Science 331, 1616–1620 (2011).

81. Babst, M., Katzmann, D. J., Estepa-Sabal, E. J., Meerloo, T. & Emr, S. D. Escrt-III: an endosome-associated heterooligomeric protein complex required for mvb sorting. Dev Cell 3, 271–282 (2002).

82. Avalos-Padilla, Y. et al. The ESCRT-III machinery participates in the production of extracellular vesicles and protein export during Plasmodium falciparum infection. PLOS Pathogens 17, e1009455 (2021).

83. Alsleben, S. & Kölling, R. Vps68 cooperates with ESCRT-III in intraluminal vesicle formation. 2022.01.03.474785 https://www.biorxiv.org/content/10.1101/2022.01.03.474785v1 (2022) doi:10.1101/2022.01.03.474785.

84. A Aguet, F., Van De Ville, D. & Unser, M. Model-based 2.5-d deconvolution for extended depth of field in brightfield microscopy. IEEE Trans Image Process 17, 1144–1153 (2008).

